# Critical period for vision-dependent modulation of postnatal retinal neurogenesis

**DOI:** 10.1101/2021.08.30.458213

**Authors:** Tatiana V. Tkatchenko, Tatyana V. Michurina, Stanislav I. Tomarev, Naoki Nakaya, Grigori N. Enikolopov, Andrei V. Tkatchenko

**Affiliations:** Department of Ophthalmology, Columbia University, New York, NY 10032, USA; Department of Pathology and Cell Biology, Columbia University, New York, NY 10032, USA; Center for Developmental Genetics, Stony Brook University, Stony Brook, NY 11794; NBIC, Moscow Institute of Physics and Technology, 123182 Moscow, Russia; Section of Retinal Ganglion Cell Biology, Laboratory of Retinal Cell and Molecular Biology, National Eye Institute, National Institutes of Health, Bethesda, MD 20892, USA

**Author notes:** **Corresponding author:** Andrei V. Tkatchenko, Edward S. Harkness Eye Institute Research Annex, Room 415, 160 Fort Washington Ave., New York, NY 10032, United States.

**Keywords:** Critical period, visual form deprivation, retinal neurogenesis, stem cells, mouse

## Abstract

It is generally accepted that retinal neurogenesis in mammals ceases shortly after birth and that stem/progenitor cells found in the postnatal eyes of mice and humans are in the quiescent state. In the present study, we have investigated postnatal retinal neurogenesis and its modulation by visual experience in the mouse model. Four age groups (P26, P45, P72, and P94) of transgenic mice expressing green fluorescent protein (GFP) in the retinal progenitor cells under the control of nestin regulatory elements were examined for the presence of nestin-GFP-positive proliferating progenitor cells in the retina. Contrary to the previously held belief, we found a significant number of proliferating progenitors at the retinal periphery in all age groups examined. The majority of these cells gave rise to photoreceptors as revealed by the genetic cell fate mapping experiments. The intensity of neurogenesis was declining with age, and strongly correlated with eye growth. Visual form deprivation resulted in a significant increase in the intensity of peripheral neurogenesis, which correlated strongly with the induced ocular growth. The susceptibility to both form-deprivation-induced increase in the peripheral neurogenesis and form-deprivation-induced increase in the ocular growth declined with age ceasing completely around P70, which marked the end of the critical period for the vision-dependent modulation of both ocular growth and postnatal retinal neurogenesis. Thus, neurogenesis in the peripheral retina of young mice is modulated by visual input, but only during a critical period in postnatal development.

## Introduction

In amphibians and teleost fish, the retina continues to grow along with the eye throughout the lifespan of the animal (Reh & Levine, 1998). Postnatal growth of the retina in these species results from the proliferation of progenitor cells located in the ciliary marginal zone of the retina (CMZ) and subsequent addition of new neurons at the retinal periphery (Johns, 1977; Straznicky & Hiscock, 1984). A population of mitotically active progenitor cells capable of differentiating into neurons was also identified in the peripheral retina of newly hatched chickens and quails (Fischer & Reh, 2000; Kubota *et al*., 2002). Unlike in the cold-blooded vertebrates and birds, it has been generally accepted that retinal neurogenesis in the mammalian eye ceases shortly after birth (Young, 1985; La Vail *et al*., 1991; Kubota *et al*., 2002) and stem/progenitor cells found in the postnatal eyes of mice and humans are in the quiescent state (Tropepe *et al*., 2000; Coles *et al*., 2004; Yip, 2014). However, we recently demonstrated that a population of proliferating neuroprogenitors persists in the retinal periphery of 3-4-month-old primates (Tkatchenko *et al*., 2006). These proliferating cells are situated both in the epithelium of the ora serrata and in the adjacent CMZ of the retina and can give rise to neurons. Furthermore, the number of proliferating progenitor cells in the retinal periphery increases significantly upon induction of experimental myopia, suggesting that ocular growth induced by form deprivation is accompanied by retinal growth. Similar to retinal growth, visual input can modulate ocular growth in cold-blooded vertebrates throughout their lifetime (Kroger & Wagner, 1996; Shen *et al*., 2005), but in warm-blooded vertebrates only at young age (Wallman & Adams, 1987; Papastergiou *et al*., 1998; Siegwart & Norton, 1998; Troilo & Nickla, 2005; Tkatchenko *et al*., 2010b). Although these studies established that visual experience can modulate ocular growth in mammals during the early postnatal period, it is not known whether this ocular growth is accompanied by retinal neurogenesis (hence retinal growth) and whether modulation of postnatal retinal neurogenesis by visual input is limited by a critical period.

In this study, we used transgenic mice expressing green fluorescent protein (GFP) in the progenitor cells of the retina (Mignone *et al*., 2004) and genetic cell fate mapping (Novak *et al*., 2000; Lagace *et al*., 2007) to study neurogenesis and the role of visual experience in proliferation and differentiation of neural progenitors in the postnatal mouse retina. Our results indicate that postnatal mouse retina harbors mitotically active progenitor cells, which differentiate into mature neurons, and that visual experience can modulate postnatal retinal neurogenesis in mice only during a well-defined critical period in postnatal development.

## Materials and methods

### Animals

C57BL/6J mice were obtained from the Jackson Laboratory (Bar Harbor, ME) and were maintained as an in-house breeding colony. Nestin-GFP transgenic mice, expressing enhanced GFP under the control of nestin regulatory elements, were maintained on the C57BL/6J background from the original stock described by Mignone et al. (2004). Z/EG (B6.Cg-Tg(CAG-Bgeo/GFP)21Lbe/J) (Stock# 004178, RRID: IMSR_JAX:004178) (Novak *et al*., 2000) and Nestin-CreER^T2^ (Stock# 016261, RRID: IMSR_JAX:016261) (Lagace *et al*., 2007) mice were obtained from the Jackson Laboratory. All animals received water and food ad libitum and all procedures adhered to the NIH guidelines for the care and use of animals in research and were approved by the Columbia University Institutional Animal Care and Use Committee.

### Analysis of progenitor cell proliferation

To label proliferating progenitor cells, P21, P40, P67, and P89 Nestin-GFP mice received ten 100 mg/kg BrdU injections over a period of 5 days (2 injections per day). Following BrdU injections, animals were sacrificed (at P26, P45, P72, and P94 respectively), eyes enucleated, posterior pole of the eyeball and the lens were removed. The “reverse” eye cups were then fixed in 2% formaldehyde in 1 X PBS for 4 hours on ice, washed in 1xPBS, cryoprotected in 30% sucrose in 1 X PBS and embedded in Tissue-Tek^®^ O.C.T compound (Sakura Finetek USA). 10-µm cryostat sections were washed three times with 1 X PBS, blocked with 5% normal goat serum, 5% BSA, 0.1% fish gelatin, 0.1% Triton X-100 and 0.05% Tween 20 in 1 X PBS for 1 h at room temperature, blocked with Mouse Ig Blocking Reagent (Vector Laboratories) for 45 min at room temperature, and then incubated with Alexa-546-conjugated anti-BrdU antibody (1:10, Invitrogen, Cat# A21308, RRID: AB_1500367), 0.1% BSA, 1 X PBS, 15 mM MgCl_2_ and 0.3 U/µl DNase I (Roche Diagnostics) for 2 h at room temperature. Sections were then washed with 0.1% Tween 20 in 1 X PBS, incubated with 300 nM DAPI in 1 X PBS, mounted in ProLong® Gold antifade reagent (Invitrogen), and BrdU/nestin-positive cells were counted under a Zeiss Axioplan 2 fluorescent microscope (Carl Zeiss Microscopy). For image capture, we used laser scanning confocal microscope Leica TCS SP5 (Leica Microsystems) and the manufacturer’s software.

### Analysis of progenitor cell differentiation

To determine whether retinal progenitor cells can give rise to neurons, P15 Z/EG/Nestin-CreER^T2^ hybrid mice received a single intraperitoneal (i.p.) injection of tamoxifen (Sigma, 200 mg/kg) diluted in corn oil (Sigma). Nine days after the tamoxifen injection (at P24), animals were sacrificed and the eye tissue sections were prepared and blocked as described above. Sections were incubated with rabbit anti-recoverin (1:2000; Millipore, Cat# AB5585, RRID: AB_2253622) and anti-GFP (1:500, Aves Labs, Cat# GFP-1020, RRID: AB_10000240) primary antibodies overnight at 4°C. The washed sections were then incubated with Alexa-594-conjugated donkey anti-rabbit (1:500, Invitrogen, Cat# A21207, RRID: AB_141637) and Alexa-488-conjugated goat anti-chicken (1:500, Invitrogen, Cat# A11039, RRID: AB_142924) secondary antibodies for 2 h at room temperature. Sections were then washed, incubated with 300 nM DAPI in 1 X PBS for 20 min at room temperature, and mounted in ProLong® Gold antifade reagent (Invitrogen). The Nestin-GFP/recoverin colocalization was examined using confocal microscopy as described above.

### Visual form deprivation and progenitor cell proliferation

To analyze the effect of changes in visual experience during early postnatal period on postnatal retinal neurogenesis, we distorted visual input by applying plastic diffusers to one of the eyes. Diffusers represented low-pass optical filters, which severely blurred the image projected onto the retina by removing high-frequency details. The diffusers were hand-made as previously described (Tkatchenko *et al*., 2010b). On the first day of the experiment (P21, P40, P67 or P89), Nestin-GFP mice were anesthetized via i.p. injection of ketamine (90 mg/kg) and xylazine (10 mg/kg), and diffusers were attached to the skin surrounding the right eye as previously described (Tkatchenko *et al*., 2010b) (the left eye served as control). Following diffuser attachment, animals received 10 i.p. 100 mg/kg BrdU injections over a period of 5 days (2 injections per day). After the last injection of BrdU, animals were sacrificed (at P26, P45, P72, and P94 respectively) and BrdU-positive progenitor cells were counted as described above.

### High-resolution MRI

MRI was done as previously described (Luan *et al*., 2006; Tkatchenko *et al*., 2010a; b). On the day of the examination, animals were anesthetized via intraperitoneal injection of ketamine (90 mg/kg) and xylazine (10 mg/kg). A contrast agent Magnevist® (117 mg/ml, Berlex Laboratories) was used as eye drops to highlight the anterior chamber of the eye. Each mouse was then positioned on an MRI-compatible holder (Bruker) and MRI data were acquired on a 7T ClinScan® MRI system (Bruker) using a two-turn transmit/receive surface coil (0.8-cm diameter, Bruker) placed over the eye. The plane for the high-resolution scan was positioned to go through the center of the lens, the center of the cornea and the optic nerve, thus ensuring its close proximity to the optical axis of the eye. Images were collected using an adiabatic spin-echo imaging sequence (repetition time, 1 second; echo time, 12 ms; number of acquisitions, 4; matrix size, 512 × 512; slice thickness, 0.60 mm; field of view, 12 × 12 mm^2^; pixel size, 23.4 × 23.4 µm^2^; 35 minutes/image). Virtual sagittal slices through the optical axis of the eye were obtained for each eye, and axial length (AL) and circumference of the eye (Circ) were measured using ImageJ version 1.44p image processing software (Abramoff *et al*., 2004). The AL was measured as a distance from the posterior surface of the cornea to the posterior surface of the retina.

### Statistical analysis

Data graphing was performed using SigmaPlot® version 10.0 (Systat Software). Power analysis was used to determine sample sizes for each experiment. Kolmogorov-Smirnov normality test was utilized to establish whether experimental data were normally distributed, and *F* test was used to check whether compared data sets had equal variances. All data sets presented here had normal distribution and equal variances, which was verified by Kolmogorov-Smirnov, Lilliefors, Shapiro-Wilk and *F* tests. We used ANOVA to analyze age-related changes in both progenitor cell proliferation and ocular growth. Paired *t* test was used to analyze differences between two repeated measures in the same experimental group, whereas *t* test for two independent samples was used to analyze differences between two experimental groups. All statistical analyses were performed using STATISTICA version 7.1 (StatSoft). All data are presented as mean ± SD.

## Results

### Postnatal retina of young mice harbors numerous proliferating progenitor cells that differentiate into neurons

We recently demonstrated that a population of proliferating neuroprogenitors persists in the retinal periphery of 3-4-month-old monkeys (Tkatchenko *et al*., 2006). To examine whether neurogenesis takes place in the postnatal mouse retina, we have analyzed mouse postnatal retina for the presence of proliferating neuroprogenitors using Nestin-GFP transgenic mice (mice expressing enhanced GFP under nestin regulatory elements) (Mignone *et al*., 2004) and BrdU as tracer to identify proliferating cells. Contrary to the previously held belief that proliferation in the mouse retina ceases shortly after birth (Young, 1985; Kubota *et al*., 2002), we found BrdU-positive proliferating cells at the retinal periphery in all age groups examined (i.e., P26, P45, P72, and P94). These cells were mostly located in the outer nuclear layers of the retina within ≈500 µm of the ora serrata, which defines retinal periphery, and expressed the neural precursor cell marker nestin as revealed by the expression of the reporter protein GFP (Figure 1).

**Fig. 1.**
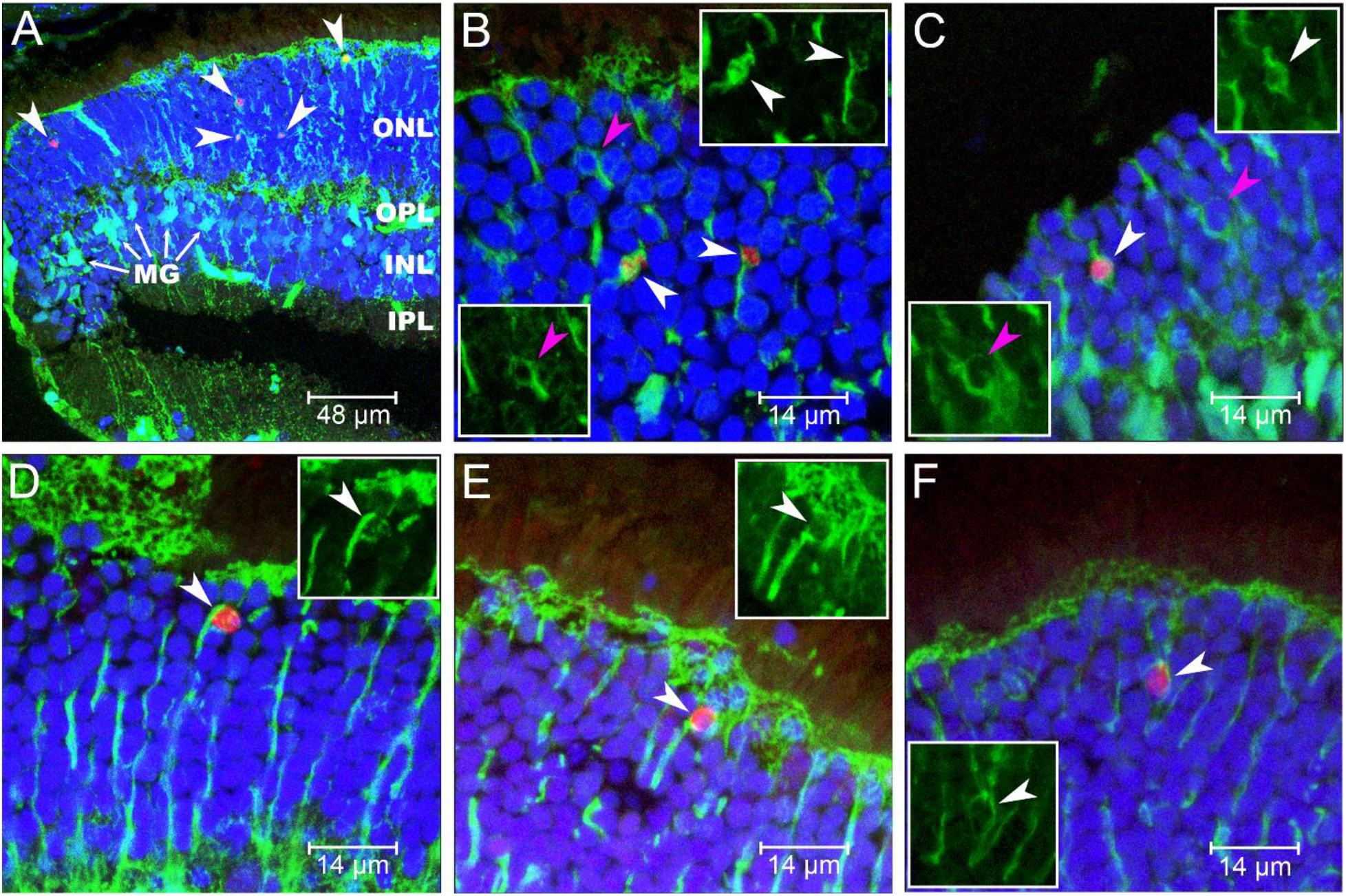
Postnatal retina of young mice has numerous proliferating progenitor cells. (A) Small magnification image showing proliferating progenitor cells at the retinal periphery of the P26 nestin-GFP mice (white arrowheads). (B-F) Large magnification images showing examples of proliferating (white arrowheads) and quiescent (magenta arrowheads) progenitor cells at the periphery of the postnatal retina of the P26 nestin-GFP mice. Green, nestin-GFP; Red, BrdU-positive nuclei; Blue, nuclei counterstained with DAPI. ONL, outer nuclear layer; OPL, outer plexiform layers; INL, inner nuclear layer; IPL, inner plexiform layer; MG, Muller glia cell bodies.

To establish whether these proliferating neuroprogenitor cells are capable of differentiating into retinal neurons, we performed progenitor cell lineage tracing analysis. Progenitor cells expressing nestin in the postnatal retina of Z/EG/Nestin-CreER^T2^ hybrid mice were irreversibly genetically labeled with GFP at P15 by inducing Cre-driven genetic recombination, which releases GFP expression in the nestin-positive cells, with a single injection of tamoxifen. The identity of the progeny of these nestin-GFP-labeled progenitor cells was than analyzed at P24. We found that the absolute majority of the postnatal retinal progenitor cells differentiate into photoreceptors, which is revealed by the double staining for GFP and a photoreceptor cell marker recoverin (Figures 2A-I). The P24 mouse retina had approximately 170 newly formed photoreceptors per millimeter of the “ciliary margin” (CMZ) (see Figure S1 for the diagram). These photoreceptors often had clearly visible developing outer segments and synaptic terminals. The differentiated photoreceptors were often found next to the undifferentiated GFP-positive/recoverin-negative progenitor cells, which had small diameter and compact nuclei suggesting asymmetrical cell division (Figures 2J-S). Thus, postnatal mouse retina harbors proliferating progenitor cells, which give rise to photoreceptors.

**Fig. 2.**
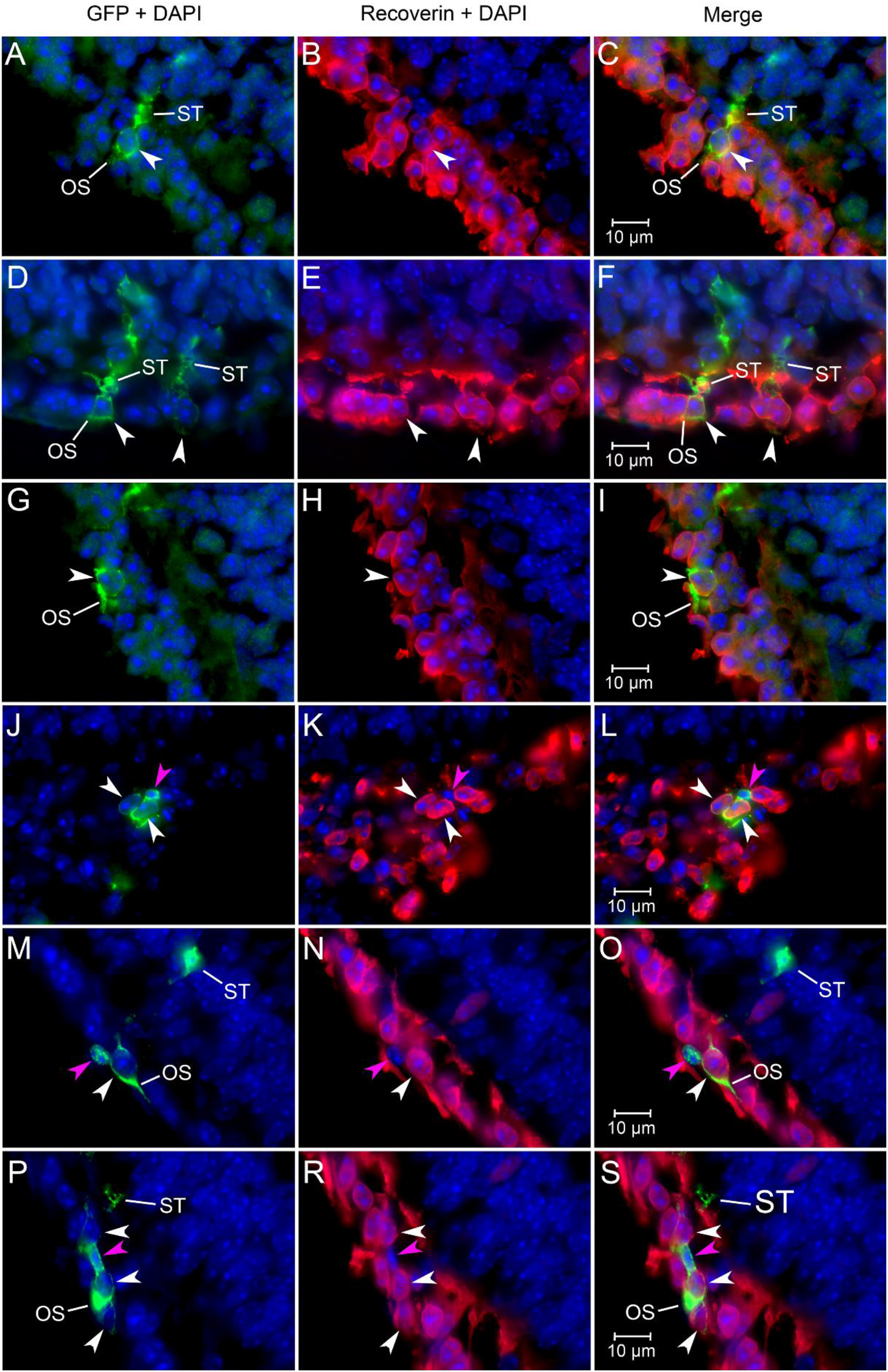
Progenitor cells in the postnatal mouse retina differentiate into photoreceptors. (A-I) Examples of progenitor cells in the postnatal retina of Z/EG/Nestin-CreER^T2^ hybrid mice genetically labeled with GFP at P15 and differentiated into photoreceptors at P24 (white arrows). Many progenitor cells formed cell clones composed of an undifferentiated progenitor cell and daughter cells differentiated into photoreceptors. (J-S) Examples of progenitor cell clones in the postnatal retina of Z/EG/Nestin-CreER^T2^ hybrid mice featuring daughter cells, which differentiated into photoreceptors (white arrows), and undifferentiated progenitor cells (magenta arrow). Double staining with antibodies to GFP (green) and recoverin (red) identifies progenitor cells, which differentiated into photoreceptors. Undifferentiated progenitors are positive for GFP, but negative for recoverin. GFP expression was released in the nestin-positive progenitor cells at P15 using tamoxifen-inducible nestin-Cre-driven genetic recombination and the identity of the GFP-labeled progenitor cell progeny was analyzed at P24, i.e., 9 days after the tamoxifen injection. Blue, nuclei counterstained with DAPI. OS, outer segment; ST, synaptic terminal.

### The number of proliferating progenitor cells in the postnatal retina declines with age

Although we found that proliferating progenitor cells persist in the mouse retina at least up to the age of three months, it was not clear whether the number of proliferating progenitors changes with age and whether there is any correlation between postnatal retinal neurogenesis and postnatal ocular growth. To answer these questions, we counted the number of proliferating nestin-GFP-positive cells in the retina of Nestin-GFP mice at different ages (Figure 3A). The retina of P26 animals had 216 ± 11 (*n* = 10) BrdU-positive/nestin-GFP-positive progenitor cells per millimeter of CMZ (proliferation index, cells/mm). The number of proliferating neuronal progenitors has exponentially decreased with age (one-way ANOVA, *F*_(3,36)_ = 2013.2, *p* < 0.0001) reaching plateau at P94 (56 ± 5 cells/mm, P45; 23 ± 4 cells/mm, P72; 16 ± 2 cells/mm, P94; *n* = 10) (Figure 3A). Interestingly, the proliferation index has reached plateau at approximately the same age as the overall ocular growth in mice, i.e., at P94 (Figure 3B); and the logarithm of the proliferation index showed a strong negative correlation with the diameter of the growing eye (*R* = -0.9958, *p* = 0.004) (Figure 3C). Thus, the intensity of neurogenesis in the postnatal peripheral retina declines with age; however, it persists into early adulthood and correlates with the ocular growth.

**Fig. 3.**
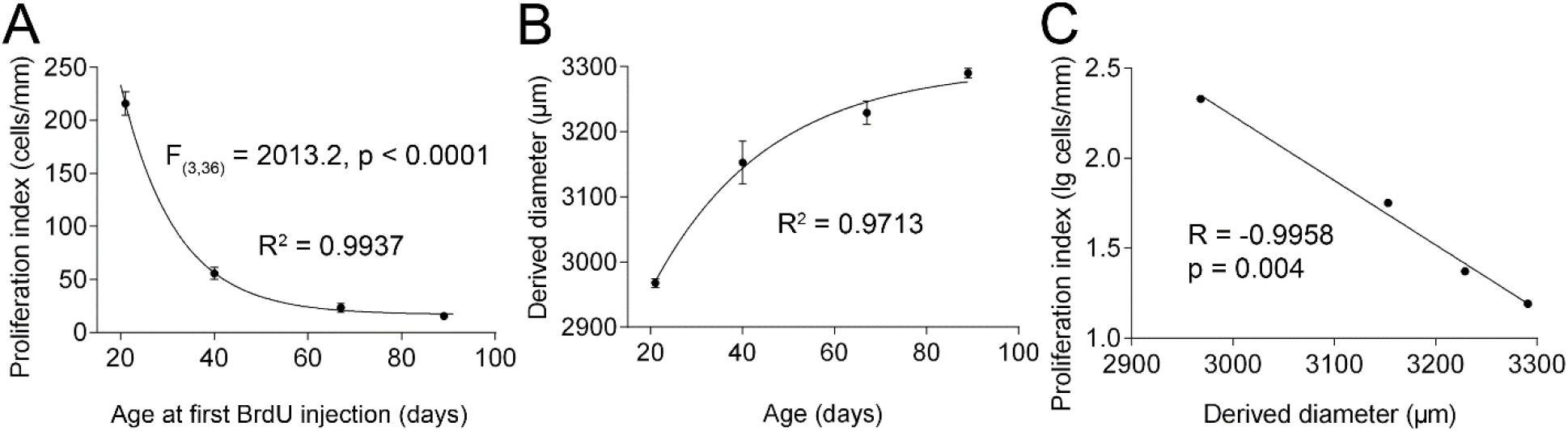
Postnatal neurogenesis in the peripheral mouse retina declines with age. (A) Analysis of progenitor cell proliferation at different ages. (B) Ocular growth in the mouse during early postnatal period. (C) The rate of ocular growth shows strong correlation with the proliferation rate of the neuronal progenitors in the peripheral mouse retina. Derived diameter is from (B) and proliferation index is from (A). Fitted lines show regressions described by the following equations: (A) y = 1194e^(−0.09x)^ + 16.93; (B) y = -757e^(−0.04x)^ + 3299; (C) y = -0.004x + 12.99. *R*^*2*^, coefficient of determination (goodness of fit); R, Pearson’s correlation coefficient; p, significance value.

### Visual experience modulates retinal neurogenesis only during a critical period in postnatal development

It was demonstrated in chickens, tree shrews and monkeys that visual form deprivation (VFD) can induce myopia (hence accelerated ocular growth) only relatively early in postnatal development (Wallman & Adams, 1987; Papastergiou *et al*., 1998; Siegwart & Norton, 1998; Troilo & Nickla, 2005). We have also recently showed that there is a susceptible period for experimentally induced myopia in mice (Tkatchenko *et al*., 2010b); however, it is not known whether vision-dependent modulation of retinal neurogenesis is confined to a critical period and whether there is any link between vision-dependent modulation of ocular growth and vision-dependent modulation of retinal neurogenesis. To investigate the role of age in vision-dependent modulation of retinal neurogenesis, we analyzed the effect of VFD on proliferation of progenitor cells in the retina of Nestin-GFP mice at different ages (Figure 4). We found that the VFD-induced increase in the progenitor cell proliferation index declined exponentially with age (one-way ANOVA, *F*_(3,36)_ = 252.07, *p* < 0.0001) (Figure 4A). At P26, we observed 91 ± 7% increase in the number of proliferating progenitors in the deprived eyes versus control (202 ± 13 cells/mm, control; 388 ± 31 cells/mm, VFD; *p* = 0.0003, *n* = 10). However, at P45, the increase was 29 ± 5% (47 ± 16 cells/mm, control; 60 ± 18 cells/mm, VFD; *p* = 0.0004, *n* = 10), and no difference was observed at P72 (34 ± 7 cells/mm, control; 34 ± 5 cells/mm, VFD; *p* = 1.0, *n* = 10). In animals analyzed at P94, we observed 8 ± 11% suppression of progenitor cell proliferation by VFD; however it was not statistically significant (30 ± 6 cells/mm, control; 28 ± 6 cells/mm, VFD; *p* = 0.18, *n* = 10). When we compared form-deprivation-induced increases in progenitor cell proliferation with the form-deprivation-induced increases in the ocular growth at different ages, we found that the logarithm of interocular difference in proliferation index showed a strong positive correlation with the interocular difference in the diameter of the eye (*R* = 0.9784, *p* = 0.02) (Figures 4B, and 4C). Importantly, the end of the susceptible period for vision-dependent modulation of retinal neurogenesis coincided with the end of the susceptible period for vision-dependent modulation of ocular growth (P72). Thus, visual input modulates retinal neurogenesis only during a critical period in postnatal development ending in mice at P72.

**Fig. 4.**
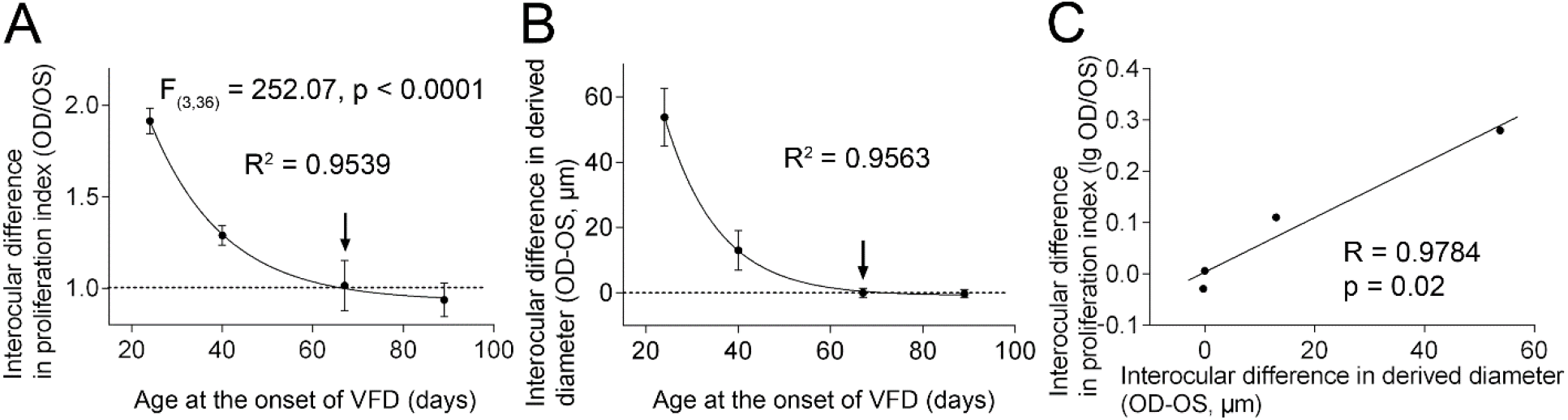
Susceptible periods for vision-dependent modulation of retinal neurogenesis and vision-dependent modulation of ocular growth in mice end at P72. (A) Analysis of form-deprivation-induced increase in retinal neurogenesis at different ages. (B) Susceptibility to form-deprivation-myopia-associated ocular growth at different ages. (C) Form-deprivation-associated increase in retinal neurogenesis is positively correlated with the form-deprivation-induced ocular growth. Fitted lines show regressions described by the following equations: (A) y = 4.36e^(−0.06x)^ + 0.93; (B) y = 432.55e^(−0.09x)^ – 0.85; (C) y = 0.005x + 0.003. Vertical arrows in (A) and (B) show the end of the critical period. R^2^, coefficient of determination (goodness of fit); R, Pearson’s correlation coefficient; OD, right eye; OS, left eye; p, significance value.

## Discussion

Several regions of the central nervous system (CNS), which are responsible for the processing of sensory inputs to the brain, undergo activity-dependent plasticity during a critical period in postnatal development (Hensch, 2004). Postnatal plasticity and critical periods have been described for the motor system (Sanes & Lichtman, 1999), somatosensory system (Erzurumlu & Gaspar, 2012), auditory system (Knudsen *et al*., 2000; Linkenhoker *et al*., 2005; Yang *et al*., 2012), taste/olfactory system (Lasiter & Kachele, 1990; Marks *et al*., 2006; Kato *et al*., 2012); however, the best studied example of a critical period plasticity is the plasticity of the visual cortex (Hubel & Wiesel, 1998; Espinosa & Stryker, 2012; Levelt & Hubener, 2012). The onset and length of the critical period for different sensory systems can vary, but postnatal plasticity is usually restricted to the early postnatal period following birth during which CNS exhibits enhanced susceptibility to both functional and anatomical remodeling. One exception from this rule is olfactory plasticity in rodents, which is not restricted to the early postnatal period, but persists throughout animal’s lifespan (Hensch, 2004). This unique situation is thought to be associated with continuous neurogenesis in the olfactory bulb, which provides the substrate for continuous remodeling of the olfactory circuits throughout the animal’s lifespan (Hensch, 2004; Lledo & Saghatelyan, 2005).

Postnatal vision-dependent plasticity has been described in the visual system at different levels of the visual pathway. The very first example of vision-dependent CNS plasticity was described in the visual cortex by Hubel and Wiesel (Wiesel & Hubel, 1963; Hubel, 1967). These authors also found that vision-dependent functional and anatomical remodeling of the visual cortex occurs only during a critical period in the early postnatal development (Hubel & Wiesel, 1970). Vision-dependent remodeling also takes place at the lower levels of the visual pathway, i.e., lateral geniculate nucleus (LGN) (Shatz, 1996) and the retina (Xu & Tian, 2007). The critical period for the vision-dependent visual system plasticity stretches from P14 (eye opening) to approximately P60 in mice (Chen & Regehr, 2000; Tagawa *et al*., 2005; Xu & Tian, 2007). Interestingly, the critical period for both visual-experience-dependent modulation of ocular growth and visual-experience-dependent modulation of retinal neurogenesis ends at approximately the same age as the critical period for the visual pathway refinement (P60-P70).

Strong correlation between ocular growth and the rate of progenitor cell proliferation in the retina during both normal postnatal eye growth and experimentally induced myopia found in this study suggests a functional link between progenitor cell proliferation and activity-dependent postnatal eye plasticity, similar to what occurs in the olfactory bulb (Hensch, 2004; Lledo & Saghatelyan, 2005). However, it has to be noted that proliferation of the progenitor cells persists in the peripheral retina somewhat longer (at least up to P94) than the vision-dependent modulation of both ocular growth and retinal neurogenesis (P72). Visual form deprivation also appears to have opposite effects on progenitor cell proliferation during the critical period and after the end of it. While form deprivation induces peripheral retinal neurogenesis in animals younger than P72, it seems to suppress neurogenesis in P94 mice, which suggests that molecular signaling required for the maintenance of progenitor cell proliferation and that required for the vision-dependent modulation of ocular growth might be distinct.

In conclusion, we found that the postnatal mouse peripheral retina has proliferating progenitor cells, which can differentiate into photoreceptors. Our data indicate that proliferation of progenitor cells in mice continues at least up to the age of 3 months, which corresponds to the age of approximately 27 months in *Macaca mulatta* and 13 years in humans based on the growth trajectory of the eye (Supplementary Table S1). These results are consistent with the finding by La Vail et al. (La Vail *et al*., 1991) that the retina of 1-month-old *Macaca mulatta* can generate new neurons and our recent finding that the postnatal retina of 3-4-month-old higher primates harbors neuronal progenitors capable of differentiating into mature neurons (Tkatchenko *et al*., 2006). Collectively, these data suggest that postnatal retinal neurogenesis is not a unique feature of the retina of low vertebrates, but is likely to be a common characteristic of the mammalian retina as well. Our data also suggest that postnatal retinal neurogenesis extends in mammals into the young adult stage of postnatal development. Moreover, peripheral retinal neurogenesis in mammals can only be modulated by vision during a critical period in postnatal development, which also coincides with the period of susceptibility for visual-experience-dependent modulation of ocular growth, which may last in humans up to the age of 14-16 years (Thorn *et al*., 2005). Strong correlation between postnatal ocular growth and the rate of neurogenesis at the retinal periphery suggests that these two processes might be connected. It would be important for the future studies to explore signaling cascades underlying the critical period for visual-experience-dependent modulation of both ocular growth and retinal neurogenesis.

## Acknowledgements

The research was supported by NIH grant R21EY018902 to A.V.T., grants from NIH (R01AG040209), NYSTEM and Russian Ministry of Education and Science (11.G34.31.0071) to G.N.E. We thank Ronald Barrett and Genene Holt for technical assistance with microscopy and the staff of the University MR Research Facility for assistance with the small animal MRI.

## Abbreviations

AL: axial length
BrdU: 5-bromo-2'-deoxyuridine
Circ: circumference of the eye
CMZ: ciliary marginal zone
CNS: central nervous system
GFP: green fluorescent protein
i.p.: intraperitoneal
LGN: lateral geniculate nucleus
MRI: magnetic resonance imaging
P: postnatal day
VFD: visual form deprivation.

